# Evaluation of Nutritional Potential and Effect of Processing on Improving Nutrient Content of Cassava (*Mannihot esculenta crantz*) Root and Leaves

**DOI:** 10.1101/2022.02.04.479097

**Authors:** Tagesse Lambebo, Tesfaye Deme

## Abstract

Cassava (*Manihot esculenta* crantz) is a major food crop in Sub-Saharan Africa, Asia, the Pacific, and South America, where it feeds around 800 million people. Its roots are an excellent source of energy, and its leaves are rich in minerals, vitamins, and protein, which could substantially supplement the current starchy diets if properly detoxified since they contain some toxic anti-nutritional factors. The objective of this study was to provide information on the nutritional profile of cassava root and leaves and role of processing in enhancing and stabilizing their valuable nutrients. Two cassava varieties, *kello* and *qulle,* obtained from Areca Agricultural Research Center were used in this study. Roots and leaves were processed (fermented, boiled, and fluid bed dried), and nutritional, anti-nutritional, and functional properties were assessed using AOAC (Association of Official Analytical Chemists) standard procedures. As compared to leaves, roots had higher carbohydrate contents, ranging from 81.0 to 87.1 g/100g, whereas leaves had higher protein (21.2 to 28.4g/100g), total carotenoid (234.1 to 987.9 μg/g), fiber (16.1 to 22.9 g/100g), Ca (499.8 to 545.4 mg/100g), K (1193.4 to 1211.0 mg/100g), Mg (274.5 to 340.5 mg/100g) and Fe (129.1 to 146.1 mg/kg) contents. Anti-nutritional factors were slightly higher in the leaves than in the roots, with toxic cyanides ranging from 10.1 to 61.0 mg/kg in the leaves and from 1.8 to 47.5 mg/kg in the roots. However, the processing of leaves decreased cyanide content from 61.0 mg/kg to 10.1 mg/kg. Similarly, other anti-nutritional factors, such as condensed tannins, oxalates, and phytate were reduced from 52.0 to 21.0 mg/100g, 156.5 to 25.9 mg/100g, and 35.9 to 4.0 mg/100g), respectively. Hence, the fermentation of leaves and roots followed by boiling and drying is an interesting approach to reduce anti-nutritional factors significantly and ensure their nutritional quality. This study revealed that variety *kello* had a relatively better nutritional profile than variety *qulle*, for both root and leaves, except for total carotenoid content, which was higher in the latter. Genetic background and processing methods can greatly affect the nutritional profile of cassava varieties, so it is critical to analyze diverse cassava germplasm and refine the processing method to identify highly nutritious and healthy varieties.

## 1. INTRODUCTION

The global food supply system has got a warning signal as a result of rapid population growth, global climate change, and health-related hazards. Countries prefer to restrict their economies, particularly the food trade, during times of crisis to guarantee that their people have enough to eat. In developing countries, malnutrition and hidden hunger are most obvious, manifesting as developmental stunting, particularly among children, and a reduced ability to withstand diseases[1]. For most African countries, reducing hunger and high food import costs is a primary objective. Countries that formerly relied heavily on food imports are looking for innovative ways to boost the use of locally grown foods. Sub-Saharan Africa (SSA) is one of the world’s poorest regions, and malnutrition remains a major issue, despite the region’s abundance of underutilized plant species (UUPS), which are high in nutrients, particularly micronutrients that are lacking in staple meals[2]. Africa is home to a wide range of nutrient-dense, underused plant species. They are excellent providers of nutrients and other health-related qualities, according to the Global Facilitation Unit for Underutilized Species (GFU), which describes them as “those plant species whose potential is not completely tapped to contribute to food security and poverty alleviation.” They have enormous promise for reducing the impact of hunger and malnutrition. The possibility of using them in key staple foods is a novel method that has a better chance of relieving poverty and malnutrition[3].

Cassava (***Manihote sculenta Crantz***) is a tuberous root crop belonging to the family ***Euphorbiaceae***, genus ***Manihot***. After rice and maize, it is the third most important food source in the tropics and the key food security crop. In the tropics and subtropics, it is a staple diet for 800 million people. It’s becoming a popular staple dish in many African countries, especially in the humid and sub-tropical tropics. It has a high nutritional content and is utilized by people in areas where food scarcity is a problem and people are malnourished. Carbohydrate, protein, vitamins, and minerals are all found in its roots and leaves[4]. It’s simple to cultivate, produces well in favorable conditions, and even produces edible roots in poor soils with dry conditions. As a result, it is drought resilient, and its mature roots can maintain their nutritional content without water for an extended period of time, suggesting that it could be the future food security crop in some developing countries[5].

According to studies, cassava can tolerate temperatures up to 40°C before the rate of photosynthesis declines. It also has a built-in system to deal with water scarcity by dropping the leaves. It can withstand salt levels of up to 150mM, with the younger ones able to withstand up to 40mM. It can live in locations where rainfall varies from 500 to 5000mm[5]. Cassava is expected to supply the world’s protein requirements. In comparison to cassava leaves, the roots have higher carbohydrate content, however the leaves and by-products of root harvest have a higher protein level (approximately 14–40%), minerals, vitamins B1, B2, and C, and carotenes. The leaves are regarded a vital protein source when compared to other green vegetables [6].

Despite their availability and nutritional benefits, incorporating these promising underutilized species into our daily diet remains a challenge. Because it includes poisonous chemicals that limit its utility, the most notable of which are cyanogenic glucosides, which are responsible for some cassava varieties’ bitter flavor. Cyanide is stored in the vacuoles of cassava cells, and it is known to be higher in the leaves and root cortex than in the root parenchyma. Several neurological disorders have been related to cyanide poisoning in areas where cassava is a major crop, including ataxic neuropathy, cretinism, and goiter and stunting in children. Cassava toxicity levels vary depending on altitude, geographic location, and the period of harvesting; crop variety and seasonal conditions. Several crops are underutilized in the majority of areas where they are grown due to a lack of awareness about their complete nutritional and anti-nutrition values, as well as information on processing. Growing drought-resistant crop varieties and employing appropriate food processing methods and equipment may help to reduce the dangers to one’s health[7]. However, the level of cyanide in the cassava can be effectively reduced with different processing methods. Processing methods such as boiling, drying and fermentation remove nearly all residual cyanogen[8]. The objective of the study was to evaluate nutritional potential and effect of processing on improving nutrient content of cassava (*Mannihot esculenta crantz)* root and leaves

## 2. MATERIALS AND METHODS

### 2.1. Samples collection

Cassava (Manihot esculenta Crantz) root and leaf samples were taken at the Areka Agricultural Research Center’s experimental farm. For this study, two types of cassava cultivars, qulle and kello, were used, and practically all experiments were carried out at Addis Ababa Science and Technology University.

### 2.2. Sample preparation

Cassava root samples were produced using the procedure outlined by [9]. To compose additional processing steps, the roots were first peeled, washed, grated, and sliced up. Weighed amounts of grated cassava pulps were soaked in deionized water at room temperature for 72hrs, by changing the water every 24hrs. After every 24hrs, the PH of the expressed juice was measured to follow the progress of fermentation. Three alternative drying techniques: sun drying, fluid bed drying and freeze drying were used in the experiment. Finally dried cassava roots were milled to a particle size of 250μm and stored in plastic bags until used in the experiment. Similarly cassava leave samples were prepared according to method described by [9]. The leaves were manually removed from the stem and washed in deionized water before being homogenized and divided into several portions for further processing (boiling, fermentation and drying). By squeezing the leaves in a sieve until the water became colorless, the samples were leached with current water for the boiling process, which took about 5min. A 700g sample mass was then weighed and placed in a Teflon-coated pan. Then 1000ml of deionized water was added, and the mixture was cooked for 15min in a single boil on a domestic cooker. To treat the samples using the fermentation method, the leaves were steeped in deionized water for 72hrs at room temperature, changing the water every 24hrs. The pH of the extracted juice was measured every 24hrs to evaluate how far the fermentation had progressed. The raw and fermented cassava leave samples were dried by sun dying for seven days, fluid bed drying at 50°C for 180min, and freeze drying for ten hours. For lowering HCN levels in cassava leaves, a temperature range of 40–80°C for 180min is beneficial [8]. To conduct the experiment, dried cassava leaves were finally ground to a powder using a crusher (laboratory mill) and packed in plastic bags.

### 2.3. Proximate composition analysis

#### ❖ Moisture Content

The moisture content of the cassava roots and leaves flour samples were determined as per AOAC[10]. About 5g of each sample was weighed into a dish, the dish and its contents were then put in an air oven maintained at 105°C and left to dry for 5hrs. The samples were then cooled in a desiccator and weighed. The dishes were then returned to the oven and dried until the moisture content variation was within 0.05%. The moisture content of the sample was then calculated as a percentage of the sample weight.

#### ❖ Ash Content

The Ash content was determined as per AOAC[10]. About 2.5g of the cassava flour was weighed in a crucible which has previously been dried in oven, cooled and weighed. Then crucible with the sample was ignited on a muffle furnace at 550°C+10°C till white ash was produced. It was cooled in desiccators for 30min. And the cooling and weighing process was repeated until the difference between two successive weighing was less than one milligram. The ash content of the sample was then calculated and expressed as a percentage of the sample weight.

#### ❖ Protein content

Crude protein content was determined as per AOAC[10]. 1g sample was weighed and transferred into a Kjeldahl digestion flask which contains 8gm of catalyst mixture (96% anhydrous (Na_2_So_4_), 3.5% copper sulfate and 0.5% selenium dioxide). Then 20ml of sulfuric acid was added and glass beads were also added to prevent bumping. The sample was first heated with maximum heating power, in order to perform the digestion within a minimum amount of time. The heating was continued until the solution become clear and the sample was then cooled down, the digestion vessel was connected directly to the nitrogen determination apparatus to carry out distillation. The product of the reaction ammonium sulfate ((NH4)2SO4) was treated by alkali (NAOH) then distilled and titrated. To the receiving flask 50ml of boric acid solution and methyl red indictors was added. The distillation apparatus with the inner condenser delivery tube dipping below the boric acid solution was connected. Then the digest was diluted with distilled water to about 30ml then neutralized with sodiumhydroxide solution and distill the ammonia into boric acid solution. Titration with sulfuric acid (0.1N) was performed after approximately 300ml has been distilled over. Finally, percentage crude protein was calculated as percentage of nitrogen.

#### ❖ Fat content

Crude fat content was determined as per AOAC [10]. 3gm well mixed dried cassava samples was weighed and place into extraction thimble for fat determination. The sample in the thimble was covered with fat free cotton wool and then the weighed flask crucible was filled with 100ml of hexane and the extraction apparatus was assembled and automatically heated for 3hrs. At the end of analysis, the extraction cups containing the extract were allowed to drying on oven (1hr at105°C) then it was cooled to room temperature in a desiccator and weighed. The percentage crude fat was calculated as percentage of sample weight.

#### ❖ Crude fiber content

Crude fiber content was determined according to AOAC [10]. 2g of the sample was accurately weighed & added into 150ml of 1.25% sulfuric acid solution in 500ml conical flask and boiled exactly for 30min after onset of boiling. The contents of the beaker were then filtered using a Buchner funnel slightly packed with previously weighed filter paper and the residue washed with hot water. The residue was then transferred into the conical flask and 150ml of 1.25% KOH solution added boiled exactly for 30min after onset of boiling. The solution was again filtered using a Buchner funnel slightly packed with filter paper. The residue was transferred quantitatively into a crucible and dried in an air oven set at 105°C for 2hrs. The desiccator was then cooled and weighed. The crucible contents were then ignited at 550°C for 3hrs and allowed to be cooled in desiccator and weighed. The crude fiber content of the sample was then calculated as percentage of the sample weight

#### ❖ Carbohydrates Content

Carbohydrates content was determined as per AOAC [10] by the difference, i.e., 100% minus the total sum of fat, moisture content, crude fiber, Ash, and proteins in 100 g of food. Percentage carbohydrate was calculated by difference.

#### ❖ Caloric value

Using the standard method of [11], the net energy value of the samples was estimated in kilo joule per hundred gram by adding up the values for carbohydrate, crude lipid, and crude protein after multiplying with their respective factors.

### 2.4. Mineral content

The mineral content was analyzed from the triple acid digestion (wet digestion method) as per AOAC[10]. About 0.5g of cassava flour was accurately weighed. 0.7ml concentrated HNO3 was added and digested on a digester for 1hour until an almost clear solution was obtained. Although no any solid particle observed, the digests were filtered using watman filter paper and volume of the solutions was made up to the mark using deionized water and introduced to the microwave plasma atomic emission spectroscopy (Agilent model MP-AES 4200) and Minerals were measured and quantifications were performed using aqueous standards for calibration. Signal response was recorded for each of the elements at their respective wavelength: Ca(393.366 nm), Fe(371.993nm), K(766.491nm), Mg(285.213nm), Na(588.995nm), and Zn(213.857nm) and the concentrations of the minerals were calculated.

### 2.5. Determination of functional properties

#### ➢ Pasting properties and mixing stability

The pasting properties of samples were determined by using Rapid Visco Analyzer (RVA), (model RVA 4500, Perten Instrunments AB, Australia) as describe by [12]. Each sample (flour weight corrected for moisture content using 3.5g at 14% moisture basis) was mixed with 25ml deionized water to obtain a final net weight (flour + water) of about 28g and placed into a canister. A paddle was then inserted and shaken through the sample before the canister was inserted into the RVA. The heating and cooling was adjusted at a constant rate of 11.25°C/min. peak viscosity, holding strength, break down, final viscosity, holding strength, set back, final viscosity, and pasting temperature were recorded with the aid of computer. The experiment was carried out within 13min in which the viscosity value was recorded every 4sec using thermocline software as the temperature increased from 50 to 95°C. The rotation speed was set to 960 rpm at the 10s initial and changed to 160rpm until the end of the experiment

#### ➢ Swelling power and solubility

The swelling power and solubility of the flours were determined using the method described in [13], with some modifications to the flour and water quantities. 1g of each flour sample was weighed into 15ml centrifuge tubes and thoroughly mixed with 10ml distilled water then mixed using vortex mixer. The mixture was heated in a water bath (Gerber Instrument, water bath, model WB-12, EU P.R.C) at 85°C for 30min, then cooled to room temperature and bench top centrifuge (Centurion Scientific, Model Pro-Analytical C2004, UK) at 3000rpm for 15min. The clear supernatant will be carefully decanted into clean, dried pre-weighed petri-dishes. The contents of the petridishes was evaporated in a hot air oven at 100°C for 30min, then removed, cooled and weighed on analytical balance. The weight of soluble substances in the supernatants was calculated by difference. The weight of each flour paste in the centrifuge tubes was also be recorded. The swelling power and the solubility of the flour samples were calculated as percentage weight.

#### ➢ Water absorption capacity (WAC)

The WAC of flour samples was determined according [13]. 1g of each flour sample was weighed into 15ml centrifuge tube and mixed with 10ml distilled water. The mixture was allowed to stand undisturbed in a test tube racket at room temperature for 30min and then centrifuged with a bench top centrifuge (Centurion Scientific, Model Pro-Analytical C2004, UK) at 3000rpm for 30min. The supernatant was carefully decanted. The weight of the flour sediment in each centrifuge tube was determined. The analysis was done in duplicate and calculated as percentage of weight

#### ➢ Titreatable acidity (TTA)

The total titreatable acidity was carried out in accordance with the method described by [14]. During fermentation period five milliliters of the sample extract were taken each 24hrs and five drops of phenolphthalein (indicator) was added to the conical flask. The mixture was titrated against 0.1N solution of sodium hydroxide. The titreatable acidity was calculated using the relation ship

#### ➢ Water activity

Water activity was determined according to AOAC [10] using (Aqua lab water activity meter, decagon instruments, Inc.) as per manufacturer’s manual.

#### ➢ Vitamin C determination

Vitamin c was determined as per AOAC [10] using iodine solution as a titrant, and starch solution as an indicator and ascorbic acid as. To prepare iodine solution (0.005M, 1.3g of iodine and 2g of potassium iodide were added into the same beaker. A few volume of distilled water was added and swirl for few minutes until iodine is dissolved. Iodine solution was transfer to 1L volumetric flask and the flask was filled up to the mark using distilled water. For the starch indicator solution (0.5%), 0.25g of soluble starch was weighed and 50ml of near boiling water was added into 100ml conical flask and stirred to dissolve and cooled before using. 1gm of cassava flour was dissolved in 100m of distilled water (in a volumetric flask) and Aliquot of the sample (20ml) solution prepared above was transferred into a 250ml conical flask, about150ml of distilled water and 1ml of starch indicator solution was added and titrated with 0.005mol L^-1^ iodine solution. The endpoint of the titration was identified as the first distinct trace of a dark blue-black color due to the starch-iodine complex. The vitamin C content was calculated from the equation:

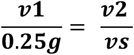

#### ➢ Total carotenoid content

Method described by [8] with some modification was used to determine total carotenoids. One gram of each grounded sample was weighed. Methanol (25ml) was then added and the mixture was transferred by vacuum filtration into a conical flask corked with filter paper. Acetone (25ml) was added to the residue to ensure maximum extraction of carotenoids. The volume of the filtrate was recorded as the volume after the first extract. The filtrate was then poured into a separating funnel (where the tap was closed) fixed to a retort stand. Petroleum ether (20ml) was placed into the separating funnel before the extract was added. Distilled water was finally added. A separation of yellowish-colored organic solvent and the colorless inorganic solvent were observed. The tap of the separating funnel was opened for the inorganic solvent to run out into a beaker placed under the funnel, and finally discarded. Distilled water was again added to the sample for washing. The inorganic solvent was collected and discarded. Washing was repeated several times until the carotenoid solution was clear. All excess distilled water was discarded. A funnel was then placed in a conical flask under the separating funnel and the organic solvent containing the carotenoids was collected. The volume of the organic solvent was recorded as the volume of the second extract. A glass cuvette was filled with the organic solvent extract and the absorbance was read at 450nm. The absorbance was read in triplicate using a UV/JASCO V-770 Spectrophotometer. TCC was then calculated using the following formula

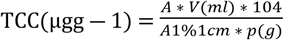

### 2.6. Determination of anti-nutrients

#### • Cyanide Content

The cyanide content of the cassava root and leave flours were determined by distillation through the alkaline titration procedure as per AOAC [10]. 10g of the grounded samples were soaked in the mixture of 200ml of distilled water and 10ml of orthophosphoric acid. The mixture was kept for 12hrs to release all the bounded cyanide. The mixture was then distilled until 150ml of the distillate was collected. To the distillate, 20ml of 0.02M sodium hydroxide was added and there after the volume of the mixture was completed up to 250ml in a volumetric flask using distilled water. The mixture was then divided into three portions, two of 100ml each and one of 50ml for both varieties and all processing parameters. 8ml of 6M ammonium solution and 2ml of 5% potassium iodide will added to 100ml aliquots, 4ml and 1ml of ammonium solution and potassium iodide was respectively added to the 50ml portions. Then it was titrated using 0.02M silver nitrate, with the 50ml portions as trials. The amount of silver nitrate consumed to the endpoint of the titration was recorded. The end point was indicated by a faint permanent turbidity appearance. The cyanide content (mg/100g cassava wet weight) of the sample was evaluated from the expression: 1mL 0.02N AgNO_3_ ≡ 1.08mg HCN.

##### Tannin content

The standard method of [15] was used with minor modification. A stock solution of 1000 ppm was prepared by dissolving 0.05g of tannic acid in 50ml of solution which consisted of 2ml of 10% sodium carbonate, 2.5ml of Folinciocalteu and 45.5ml of 70% acetone. By using the relation M1V1=M2V2 (where M1 and M2 represent concentrations; V1 and V2 represent volume), concentration of 1.0, 2.0, 3.0, 4.0 and 5.0ppm was prepared from the stock solution of 1000ppm. For cassava flour, 0.5g of the sample was weighed into a 100ml bottle, and then 50ml distilled water was then added and shaken very well for an hour using a mechanical shaker. The solution was filtered into a 50ml volumetric flask and made up to the mark. The 5ml of the filtrate was transferred into a test tube and mixed with 2ml of 0.1M FeCl3 in 0.1NHCl and 0.008M potassium Ferro cyanide. The absorbance of these tannic acid concentrations was measured at 725nm using a UV/JASCO V-770 Spectrophotometer). A regression equation was obtained by plotting a graph of absorbance against the concentrations which was used to determine the tannic acid content of each sample extract. The total tannic acid was expressed as mg TA equivalent/100g of sample

#### • Phytate Content

The standard method of [16] was used for tannin determination. 4g of the cassava flour was soaked in 100ml of 2% HCl acid for 3hrs and filtered. Then 25ml of the filtrate, 5ml of 0.3% NH4SCN and 53ml of distilled water were mixed together and titrated against 0.01M standard FeCl3 until a brownish yellow color persist for 5sec. Phytin phosphorus (1ml = 1.19mg phytin phosphorous) was determined and the phytic acid content was calculated by multiplying the value of the phytin phosphorus by 3.55

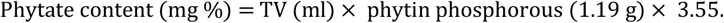

Where TV is the volume obtained after color change.

#### • Oxalate content

Oxalate content was used to determined standard method of [17]. 2g of the flour residue was weighed into a 250ml volumetric flask containing 190ml of distilled water and 10ml of 6M HCl. The mixture was digested for 1hr in a boiling water bath, cooled and made up to the mark and then filtered. The 50ml aliquot of the sample was measured into a beaker and 20ml of 6M HCl was added. The mixture was evaporated to about half of the volume and then filtered. The residue was then washed several times with warm distilled water and 3 drops of methyl orange indicator was added to 25ml of the filtrate and titrated against 0.05M KMnO4 solution till a faint pink color appeared and persisted for 30sec. The total oxalate content was finally calculated

### 2.7. Statistical Analysis

Three processing techniques and two varieties were used in the factorial designs (3×2 factors). However, the drying procedure was used in three different setups, and this was used as a test to see which one was best for nutrient retention and detoxification. The data obtained from the study was presented in a tabular form. SPSS Version 26 was used for all calculations. The mean values for samples with significant differences were compared using a two-way ANOVA with the Tukey multiple-range test at p<0.05.

## RESULT AND DISCUSSION

### 2.8. Proximate Composition of the cassava root and leaves

Proximate compositions of cassava root were presented in table 1, thus, the following concentration ranges were obtained for cassava root samples: moisture (5.85to 7.30%), ashes (0.8to 2.4%), proteins (0.25 to 1.25%), fat (2.01 to 3.70%), fiber (0.98 to 2.31%) carbohydrates (81 to 87.08%) and caloric value (347.25 to 378.33%). Whereas, the proximate compositions of cassava leave samples were presented following concentration ranges between: moisture (6.11 to 7.79%), ashes (6.22 to 9.81%), proteins (21.25 to 28.45%), fat (4.86 to 7.79%), fiber (16.15 to 22.95%) carbohydrates (30.62 to 39.16%) and caloric value (269.18 to 311.85%).

**Table 1:**
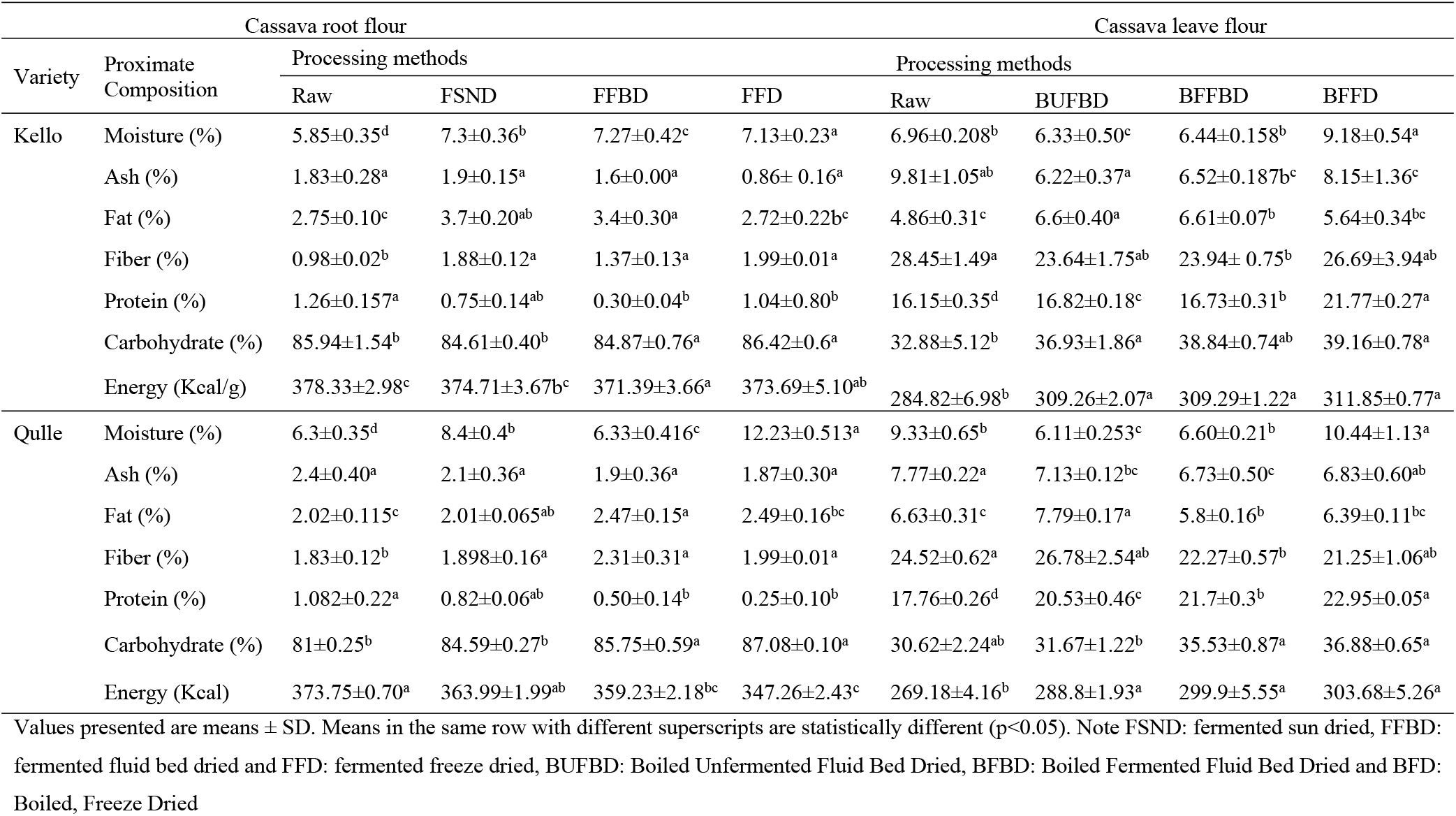
Effect of variety and the processing methods on proximate composition of cassava root and leave flour.

Moisture content was significantly (p < 0.05) affected among all processing conditions and both varieties. According to report of [12] moisture content of cassava root flour raged between 5.97 to 6.33%. The observed values in this study were comparable with that reported in the literature. But, there was a small difference because of variations in variety of cassava samples and the method of drying as well as the moisture content of the dried cassava roots before it was milled into flour. Whereas the moisture content of the cassava leaves flour in this study was in the range between 6.11 and 7.79%. The maximum moisture content (7.79%) was observed is BFFD qulle variety and minimum value (6.11%) for BUFFBD in kello variety. There was a significant difference (p<0.05) in moisture contents in both varieties and among processing methods. The moisture content in the present study was slightly lower than that reported by [18], in which the moisture content for selected local varieties of cassava leaves was ranged between 21.5 and 36.4%. The small difference in moisture content of cassava leaves may be because of the difference in varieties and the method of processing. The moisture content of the present study (6.11 to 7.79%) was within the range that reported by [19] in which the values were in the range between 5.05% and 11.6%. This indicates that low moisture content of cassava leave flour as important parameter to prolong shelf life.

Ash content of cassava root flour in this study was between 0.8% and 2.4%. The maximum value (2.4%) was observed in FFBD kello variety and the minimum value (2.01%) for qulle variety and the lowest value (0.8%) was observed in FFD qulle variety. The ash content was significantly different (p<0.05) for both varieties. The difference in the ash content may be as a result of the varietal difference. The values recorded in present study (0.8% and 2.4%) were comparable to results obtained by [20], in which the ash contents of was between 1.21 and 1.78% and by [20] in which ash content ranged between 2.19% and 2.43%. This suggests that cassava root varieties in this study contain significant amount of minerals. The recorded values were slightly lower than that reported by [17] for the same cassava root varieties in between 2.43% and 3.45% Kello and Qulle respectively. This slight difference might be as a result of several intrinsic and extrinsic factors such as age of cassava varieties, soil type, fertilizers and production conditions. Whereas, the ash content of cassava leaves in present study was in the range between 6.22 and 9.81%. It was significantly (p<0.05) affected by both variety and processing methods. The maximum ash value was observed in raw kello variety and the content was markedly decreased as its fermentation and thermal processing. According to [21]. Ash contents of different cassava leave varieties were in the range between 4.1 and 6.16%. And [19] reports, the ash content of cassava leaves were in the range between 5.68 and 6.13%. The results obtained in present study (6.22 to 9.81) was within the range of findings reported by [22] in which the average ash content of cassava leave ranged between 7.36 and 10.5%. This indicates that cassava leaves used in this study contain significant amount of minerals. A decrease in ash content during processing might be associated with leaching of some minerals into the water. Relatively high ash content in cassava leaves than roots indicate high mineral content. A high mineral element in food enhances growth and development and also catalyzes metabolic processes in human body.

Fat content in cassava root flour was ranged between 2.01% and 3.7%. The maximum value (3.7%) was observed for FSND and the minimum value 2.01%) for FFBD qulle variety. The fat content was significantly different (p<0.05) for both varieties and in all processing parameters. According to report [23] ash content of cassava root flour ranged between 0.55 to 0.69% [20] reported that the range of crude fat contents in between 0.15–0.63%. This slight difference might be as a result of several intrinsic and extrinsic factors such as age of cassava varieties, soil type, fertilizers and production conditions. Whereas, the fat content of cassava leaves flour in this study was in the range between 4.86 and 7.79%. The values were significantly (p>0.05) affected by processing methods and varieties as well. The results (4.86 and 7.79%) were in the range between the results that was reported by [21] in which fat content of cassava leave varieties were in the range between 3.84 and 9.1%. The values were slightly greater than fat content (1.01 and 4.81%) reported by [24]. The slight difference might be due to its difference in variety, processing methods, age and agro ecological factors of cassava leaves.

Fiber content of cassava root flour in this study was ranged between 0.98 and 2.31%. The maximum value (2.31%) was recorded for FFBD qulle variety and the minimum amount (0.98%) was recorded for raw kello variety. The fiber content was significantly different (p<0.05) for both varieties and among all processing methods as well as for the interaction of variety and processing. The crude fiber content of cassava roots was reported between 3.32 and 3.78 % by [20]. [25] also reported that fiber content ranged from 0.03 to 0.60%. The crude fiber content of cassava root flour ranged from 2.09 to 2.38% according to a report [12]. The current experimental value (0.98-2.31%) was in the middle of the above-mentioned results. This indicates that cassava root varieties in this study contain significant amount of dietary fiber. Whereas, the crude fiber content of cassava leave in this study was ranged between 16.15 and 22.95%. For both kello and qulle varieties, the crude fiber content was significantly (p<0.05) affected by all processing conditions. The maximum fiber content was observed for FFD qulle variety. This indicates that the crude fiber contents of cassava varieties can significantly increase by processing especially by fermentation. The experimental value was slightly greater than the fiber content of some cassava leave varieties reported by [22]: 10.09 to 16.10%, [21]: 9.16 to 15.61% and [25]: 11.60 to 14.28% and [19]: 18.6 to 19.5%. This slight difference might be due the difference in variety, agro ecological factors, maturity and processing parameters.

Protein content of cassava root flour in this study was between 0.25% and 1.26%. The maximum value 1.26% recorded for raw kello variety and minimum (0.25%) for FFD qulle variety. The processing methods had a significant (p<0.05) effect on protein content, whereas the interaction of both variety and processing had no significant (p>0.05) effect on protein content of cassava root flour. According to reports [8] the protein content of cassava root flour ranged between 0.01 and 1.45% and that reported by [20] in the range between 1.21 and 1.87%. The values in present study (0.25% to 1.26%) lie between the values reported in literature. But the protein content in present study was slightly lower than that reported by [23] in which the protein content of cassava root was between 2.09 and 2.69%. This slight difference may be due to intrinsic factors such as variety and maturity. Whereas, the protein contents of Cassava leaves in present study were varied from 21.25 to 28.45%. These values were similar to the values observed for cassava leaves by [26]: 21.6 to 32.5%, [18]: 21.5 to 36%, [22]:17.13 to 36.93 [24]: 19.73 to 29.47% and [21]: 21.51 to 28.73%. The protein content of cassava leaves were significantly (p<0.05) affected by both varieties and all processing parameters. The variation in protein content could be due to variations in leaching of some soluble proteins and the degree of denaturation of other groups of proteins following different drying processes. The results in this study lies within the result reported by [19], in which the protein content of cassava leave was (17.7-38.1%) which indicates that cassava leave is a good source of protein.

Carbohydrate content of cassava root flour in this study was ranged between 81.0 to 87.08%. It was significantly (p<0.05) affected all processing parameters and both varieties. There was an increase in the amount of total carbohydrates when comparing raw to processed cassava root and the highest carbohydrate content (87.08%) was observed for FFD qulle variety. The increase in carbohydrate content after the drying and fermentation process may be due to the fact that some high molecular weight carbohydrates such as starch are broken down into less complex sugars other byproducts [20] reported that the range of carbohydrates content for cassava root flour was between 84.32-86.57%. The present experimental value (81 to 87.08%) lies between the reported values by [27] in which the total carbohydrate content of cassava root was between 83.86-91.33%. This indicated that varieties and processing methods in this study could retain significant amount of carbohydrates in cassava root. Whereas, the carbohydrate content for cassava leaves in this study was ranged between 30.62 and 39.16%. It was significantly (p<0.05) affected by varieties and processing techniques. The carbohydrate content was relatively greater in kello variety than that in qulle variety. The content was increased during processing this might be due to the breakdown of higher molecular weight carbohydrates into less complex sugars and resulting in the release of sugar from the complex matrix. According to [18] total carbohydrate content of cassava leave ranges between 47.2 to 68.1%. As reported by [24], the carbohydrates content of cassava leaves varied from 66.1 to 72.1%. The results in this study (30.62 to 36.16%) was close to that reported by [21] in which the carbohydrate content of different cassava leave were reported between 35.17 and 48.80%.

Total caloric value of cassava root was ranged between 347.25 and 378.33. The values were significantly different (p<0.05) for both varieties and for all treatments as well as the interaction effect was also significant. Whereas, the caloric values for cassava leaves in this study were in the range between 269.18 and 311.85%. In both varieties the caloric value was significantly (p>0.05) affected by processing methods. the values in present study was slightly lower than findings reported by [24], in which total caloric values for selected varieties of cassava leaves were between 395 and 425kcal/g. The slight differences might be due to varietal difference, soil type, and agro ecological factors.

### 2.9. Mineral compositions of cassava root and leave

Table 2, shows the mineral composition in two local varieties of cassava. Thus, the following concentration ranges were obtained for both root and leave samples in this study: Ca(52.49-54.90mg/100g), Mg(38.25-57.74mg/100g), K(292.38-296.77mg/100g), Na(219.39-239.79ppm), Fe(14.28-16.65ppm) and Zn(15.33-18.61ppm) for cassava root and Ca(499.85-545.36mg/100g), Mg(274.47-340.53mg/100g), K(1193.37-340.53mg/100g), Fe(89.36-121.96mg/100g), Na (129.14-146.12ppm) and Zn (119.46-132.11ppm) for cassava leave with the order of: K> Ca > Mg> Na>Fe> Zn. The mineral composition of both varieties of root in this investigation revealed significant differences (p<0.05). The variations might be due to the mineral content of the soil from which the plants absorb their nutrients. The values were similar to those found by [28] in which the mineral concentrations of cassava root were Ca (22.4-238.6 mg/100g), Mg (24.52-165.22 mg/100g), Fe (75.38-606.37ppm), and K (22.42-22.44 mg/100g) and higher than that reported by [27] in which the content was between Ca(13.15-18.09mg/100g), Fe (0.00-0.01mg/100g), Zn (0.56-0.87mg/100g), and Mg (0.56-0.87mg/100g) (3.58-3.88). The values in this study were comparable with that reported by [29] in which the mineral contents were in the range between Ca(79.196-115.284mg/100g), Na(25.58-28.25ppm), K(69.28-76.41mg/100g), Fe(412.70-422.48ppm), Mg(31.76-33.34mg/100g), and Zn(17.52-22.46ppm). While Several cassava leave varieties showed mineral concentrations in the range between Ca(31.1-1360mg/100g), K(27.08-1870mg/100g), Mg(15.77-488.78mg/100g) and Fe(15.77-488.78ppm) and Zn(20.2-542.71ppm) as per [28]. According to [30], cassava leaves contains Ca (680mg/100g), Mg (20-70mg/100g), K (120-160mg/100g), Na (32.78-33.45ppm), Fe (321.32-458.08ppm) and Zn (321.32-458.08ppm) (36.25-68.95ppm). [19] in their investigation reported the mineral content of cassava leaves as K(1380-2260 mg/100g), Ca(430-1140 mg/100g), Mg(260-370 mg/100g), Na(38-120 mg/100g), Fe(15-27 mg/100g), and Zn(12-21 mg/100g). This suggests that the cassava leaves in this study have a comparable mineral makeup and it could be used to supplement other food recipes. The kello variety had relatively higher levels of mineral content except Na. This indicates that the kello variant was relatively superior in terms of mineral composition.

**Table 2:** Mineral contents of two different cassava cultivars.

| Contents | Cassava root varieties |  | cassava leaves varieties |  | V |
| --- | --- | --- | --- | --- | --- |
|  | Kello | Qulle | Kello | Qulle | ** |
| Na (ppm) | 219.39±0.84 | 239.79±1.36 | 129.14±0.62 | 146.12±3.46 | ** |
| Ca (mg/100g) | 54.90±9.31 | 52.49±2.80 | 545.36±9.51 | 499.85±21.500 | ** |
| K (mg/100g) | 296.77 ±4.33 | 292.38±5.54 | 1211.03±4.10 | 1193.37±7.24 | ** |
| Fe (ppm) | 16.65±9.71 | 14.28±0.97 | 121.96±13.83 | 89.36±16.48 | ** |
| Zn (ppm) | 18.61±2.76 | 15.33±7.95 | 132.11±2.23 | 119.46±0.55 | ** |
| Mg (mg/100g) | 57.74±0.18 | 38.25±0.73 | 340.53±2.32 | 274.47±2.66 | ** |

Values presented are means ± SD. Note V: variety and \*\* stands for significant difference.

### 2.10. PH and Tititratable acidity (TTA) of cassava root and leaves

Titiratable acidity and pH are two interrelated concepts in food analysis that deal with acidity. The capacity of microorganisms to grow in a certain food can be determined using its PH, whereas titreatable acidity is more accurate in predicting how organic acids in the diet affect flavor [33]. The PH and titreatable acidity of cassava root and leaves throughout fermentation was illustrated in table 3. PH of the cassava root in this study was dropped rapidly during fermentation from 7.25 to 4.4 and 6.65 to 4.46 for kello and qulle varieties respectively. The rapid drop in pH during cassava root fermentation was comparable to that previously reported by [31], in which the PH of cassava root dropped from 6.34 to 4.21during 48hrs of fermentation. There was a gradual increase in total acidity during fermentation from 0.041 to 1.8% and 0.06 to 2.31% for kello and qulle varieties respectively. The observed changes in PH and TTA during cassava root fermentation in this study were probably due to the accumulation of organic acids. The results from this study might ensure that cassava root used in present study undergone fermentation to improve nutritional value and to reduce anti-nutritional factors to a safe level of consumption. Whereas, PH of cassava leaves did not drop as that of cassava roots in this study because, cassava leaves had a lower concentration of carbohydrates or starch to be breakdown during fermentation. In this study, the pH of cassava leaves was increased from 5.96 to 7.41 and 6.29 to 7.06 for kello and qulle varieties respectively. The TTA of the kello and qulle varieties were decreased slightly during fermentation, from 0.17 to 0.010% and.108 to 0.0230%, respectively. The values in this study were close to the result previously reported by [18] in which the range was between: 0.16 to 1.17% for titreatable acidity and 5.91 to 6.41 for PH. The PH of cassava leaves increased from 6.5 to 8.5 during fermentation, according to the report [32], and the value of titiratable acidity dropped as alkaline compounds were formed during fermentation. The slightly alkaline PH recorded during cassava leave fermentation might be attributed to amines produced by bacillus. Bacillus strains can use cyanohydrin as a source of nutrition, particularly Bacillus pumilus. As a result, they could be able to help with cyanide reduction in the fermentation medium[32].

**Table 3:** pH and Tititratable acidity (TTA) as affected by fermentation.

| Cassava roots |  |  |  |  |  |  | Cassava leaves |  |  |  |  |  |
| --- | --- | --- | --- | --- | --- | --- | --- | --- | --- | --- | --- | --- |
| Variety | PH |  |  | TTA (%) |  |  | PH |  |  | TTA (%) |  |  |
|  | Day1 | Day2 | Day3 | Day1 | Day2 | Day3 | Day1 | Day2 | Day3 | Day1 | Day2 | Day3 |
| Kello | 7.25±0.35 | 4.98±0.01 | 4.4±0.1 | 0.041±0.01 | 1.59±0.43 | 1.8±0.2 | 5.96±0.01 | 6.14±0.03 | 7.41±0.2 | 0.17±0.01 | 0.02±0.05 | 0.01±0.00 |
| Qulle | 6.65±0.15 | 4.95±0.01 | 4.46±0.16 | 0.07±0.01 | 2.18±0.38 | 2.31±0.31 | 6.29±0.02 | 6.63±0.01 | 7.06±0.2 | 0.11±0.00 | 0.06±0.01 | 0.02±0.00 |

### 2.11. Functional properties of cassava root and leaves

Water absorption capacity describes flour–water association ability under limited water supply. It depends up on the power of aggregation between starch molecules. Weak aggregation power between starch molecules causes the surface of its molecules to form a bond with water molecules become easier thus increase the rate of water absorption capacity [12]. The water absorbing capacity of cassava root flour was presented in table 4, and the values observed were in the range between 288.5 and 474% and the values were significantly (p<0.05) affected by all processing methods. The differences in WAC might likely be due to difference in types of cassava root four and process incorporated. Whereas, the water absorbing capacity of cassava leave flour presented in this study was between 211 and 546%. These values were significantly (p<0.05) affected by processing methods and varieties. The WAC of the cassava leaves in the present study were much higher than the values reported by [33] in which water absorption capacity of cassava leaves flour were in the range between 404 and 417%. These variations might be due to difference in processing and analyzing methods. The higher WAC values in this study indicate that cassava leaves might be suitable in the formulation of foods.

**Table 4:** Functional properties under different processing conditions in cassava root flour.

|  |  | Cassava root |  |  |  | Cassava leaves |  |  |  |
| --- | --- | --- | --- | --- | --- | --- | --- | --- | --- |
|  |  | Processing methods |  |  |  | Processing methods |  |  |  |
| Variety | Parameters | RAW | FSND | FFBD | FFD | RAW | BUFBD | BFFBD | BFFD |
| Kello | WAC (%) | 474±9.89<sup>a</sup> | 338.5±14.8<sup>b</sup> | 410±8.48<sup>b</sup> | 378±15.55<sup>c</sup> | 546±29.69<sup>a</sup> | 410.5±10.60<sup>b</sup> | 355.5±10.60<sup>c</sup> | 284.5±6.36<sup>d</sup> |
|  | SWP (g/g) | 6.54±0.39<sup>a</sup> | 6.02±0.02<sup>b</sup> | 5.29±0.12<sup>a</sup> | 4.44±0.06<sup>c</sup> | 4.81±0.148<sup>a</sup> | 2.69±0.17<sup>b</sup> | 3.505±0.16<sup>b</sup> | 2.015±0.26<sup>c</sup> |
|  | Solubility (%) | 2.71±1.00<sup>a</sup> | 2.55±0.7<sup>ab</sup> | 2.42±0.57<sup>ab</sup> | 1.43±0.60<sup>b</sup> | 6.61±3.12<sup>a</sup> | 7.24±0.44<sup>a</sup> | 9.5±0.38<sup>a</sup> | 6.36±0.69<sup>a</sup> |
|  | Aw | 0.46±0.01<sup>b</sup> | 0.45±0.01<sup>b</sup> | 0.47±0.01<sup>c</sup> | 0.67±0.01<sup>a</sup> | 0.60±0.00<sup>a</sup> | 0.55±0.02<sup>b</sup> | 0.52±0.01<sup>c</sup> | 0.68±0.00<sup>d</sup> |
| Qulle | WAC (%) | 400±2.83<sup>a</sup> | 442.5±12.02<sup>b</sup> | 399.5±2.12<sup>b</sup> | 288.5±13.43<sup>c</sup> | 446±29.69<sup>a</sup> | 314±14.14<sup>b</sup> | 270.5±38.89<sup>c</sup> | 211±12.73<sup>d</sup> |
|  | SWP (g/g) | 6.22±0.04<sup>a</sup> | 5.165±0.20<sup>b</sup> | 4.87±0.07<sup>a</sup> | 4.26±0.34<sup>c</sup> | 3.74±0.13<sup>a</sup> | 3.12±0.31<sup>b</sup> | 2.615±0.33<sup>b</sup> | 1.5±0.59<sup>c</sup> |
|  | Solubility (%) | 3.72±0.40<sup>a</sup> | 1.04±0.06<sup>ab</sup> | 2.19±0.26<sup>ab</sup> | 1.76±0.06<sup>b</sup> | 11.75±1.34<sup>a</sup> | 11.13±3.01<sup>a</sup> | 9.75±1.77<sup>a</sup> | 9.83±0.25<sup>a</sup> |
|  | Aw | 0.41±0.00<sup>b</sup> | 0.45±0.020<sup>b</sup> | 0.33±0.01<sup>c</sup> | 0.50±0.02<sup>a</sup> | 0.55±0.01<sup>a</sup> | 0.51±0.01<sup>b</sup> | 0.42±0.01<sup>c</sup> | 0.68±0.01<sup>d</sup> |

Values are means ± SD; Means not sharing a common superscript letter across the rows were significantly different (P≤0.05)
Note: FSND: Fermented Sundried, FFBD: Fermented Fluid Bed Dried, FFD: Fermented Freeze Dried, BUFBD: Boiled Unfermented Fluid Bed Dried, B FBD: boiled Fermented Fluid Bed Dried and BFFD: Boiled fermented and Freeze Dried, WAC: Water Holding Capacity, SWP: Swelling Power and aw: water activity

Swelling power and water solubility decreased along with increasing fermentation time caused by lack of nutrients [34]. The solubility of the cassava root flour was in the range between 1.04 and 3.72%. The solubility values in this study was not significantly (p>0.05) affected by varieties while the values were significantly (p<0.05) by processing parameters and interactions. The present result 1.04 to 3.72% was lower than the results presented by [35] in which the solubility of cassava root flour was in between 13.3 and 18.18%. The observed variations may be due to differences in types of processing. Whereas, the solubility of the cassava leaves flour was ranged from 6.36 and 11.75%. There was no significant difference (p>0.05) between varieties while results were significantly different for processing parameters and for the interaction effect. The present result 6.36 and 11.75% was lower than the results presented by [35] in which the solubility of cassava root flour was in the range between 13.3 and 18.18%. The observed variations may be due to processing parameters.

Heating starch molecules in excess water results in disruption of crystalline structure and exposure of hydroxyl group. The formation of hydrogen bond by hydroxyl group and water molecules results in the swelling and solubility of starch granule[36]. Swelling power provides evidence of non-covalent bonding between starch molecules and is influenced mainly by the ratio of amylose to amylopectin [37]. The swelling power of the cassava root flour in this study was ranged from 4.25 to 6.53g/g. The values recorded were significantly (P<0.05) affected by both variety and processing conditions. The values in this study were lower than that reported by [35] in which the swelling power of cassava root flour was reported between 21.7 to 25.37g/g. The presence of non-starch components such as proteins and lipids in different varieties and increased amylase activity during processing such as fermentation might be responsible for difference in swelling power [20]. Whereas, the swelling power of the cassava leave flour in this study was between 1.5 and 4.8g/g. The values recorded were significantly (P<0.05) affected by both variety and processing conditions. And the higher solubility and lower swelling power in cassava leave flour could be important for different food formulations.

Water activity is a measure of the energy status of the water in a system and thus is a far better indicator of perishability than water content [38]. The content of free water used by microorganisms for growth is referred to as water activity. It’s linked to feed moisture content (high feed moisture content implies high water activity values, and vice versa) [39]. Growth of fungi and some yeast usually require water activity between 0.6 and 0.7. The water activity values of cassava root flour were presented in table 4, in the range between (0.33 and 0.67). These lower values indicate that cassava root flour is shelf stable with respect to water activity value. Whereas, water activity of cassava leaves presented in table 9, were in the range between 0.33 and 0.50%. The values were significantly (P<0.05) affected by both varieties and processing methods. Low water activity levels (0.33 to 0.50) were detected for all processing parameters. The water activity of cassava leave flour in this study (0.33 to 0.50) was close to that reported by [7], who reported that, the water activity of cassava leaf flour was between 0.38 and 0.43. This suggests that cassava leaves flour may have the best storage stability in relation to water activity.

Peak viscosity (PV) is an important rheological property of starchy foods, and reflects the behavior of flour and starch paste under various shears, temperature and time [40]. Peak viscosity is the ability of starch to swell freely before their physical breakdown and measures the maximum viscosity developed during or soon after heating. High peak viscosity is an indication of high starch content and also relate to the water-binding capacity of starch, Samson [41]. Flour with a lower peak viscosity has a lower thickening power than flour with higher peak viscosity [42]. PV in this study decreased significantly (P<0.05) from 7832cp in raw qulle variety to 4875cp in FFD qulle kello variety. This difference could be related to starch molecule breakdown during fermentation, plant maturity, harvest season, paste concentration and compositional differences in species.

Peak time is the corresponding time required for viscosity to become peak (maximum). The values of Pt in this investigation ranged from 6.9 minutes in raw qulle to 9.15 minutes in raw kello and the mean peak time of cassava root flour from two cassava variants was 8.59 min, however they were not significantly different (p>0.05) among varieties while they were significantly different (P<0.05) throughout processes. The average peak time in this study was slightly greater than that reported by [43] in which the average peak time was 4.98 minute.

Trough viscosity (TV) and breakdown viscosity (BV) are measurements of resistance and disintegration of flour or starch respectively, when exposed to temperature changes during pasting. In this study, the TV of cassava root flour increased significantly from 1583cp in raw kello to 2505cp in FFBD qulle varieties, and it was significantly different (P≤0.05) between varieties and processing conditions. The value of TV in present study was slightly lower than that reported by [37] in which the average values of TV of cassava was in between 2724 to 3369cp. The difference in TV might be due to the difference in varieties and age of cassava samples.

Breakdown viscosity measures the susceptibility of the starch granule to disintegrate (degree of viscosity reduction) during heat processing and this may affect the stability of the flour product [44]. High breakdown viscosity is a disadvantage in many food applications because it results in unevenly distributed viscosity [42]. It can be used as an indicator for pasting stability during the heating and stirring. The qulle variety FFBD flour had the highest breakdown viscosity (5233cP), while raw kello variety flour had the lowest one (3212.5cP). BV in this study was significantly (P<0.05) affected by processing. This indicates that a major modification occurred in starch granules of cassava root after processing methods such as fermentation and different drying techniques. This finding indicates that, qulle variety flour had the ability to withstand severe processing conditions better than kello variety. The BV value was also highest in the qulle FFBD flours than all other flours (Table 5). This suggest that, the swollen starch granules of fermented and fluid bed dried flours can be easily been disintegrated than those of raw and sundried starch granules. Thus, unfermented starch granules are more stable during heating process than fermented starch granules.

**Table 5:** Pasting properties of cassava roots as affected by varieties and processing methods.

| Variety | Parameters | processing methods |  |  |  |
| --- | --- | --- | --- | --- | --- |
|  |  | Raw | FSND | FFBD | FFD |
| Kello | PV (cp) | 7270.5±31.11<sup>b</sup> | 7158±22.63<sup>a</sup> | 4875±31.11<sup>d</sup> | 5634±155.56<sup>c</sup> |
|  | TV (cp) | 1583.5±48.79<sup>b</sup> | 2316±7.07<sup>a</sup> | 2397±101.8<sup>a</sup> | 2129.5±125.16<sup>a</sup> |
|  | FV (cp) | 2559±120.2<sup>c</sup> | 2521.5±74.24<sup>b</sup> | 3491.5±45.96<sup>a</sup> | 3402±18.38<sup>a</sup> |
|  | BV (cp) | 3212.5±94.05<sup>b</sup> | 4952±60.81<sup>a</sup> | 4692.5±221.3<sup>a</sup> | 3386.5±197.28<sup>b</sup> |
|  | SV (cp) | 258.5±6.36<sup>d</sup> | 1001±35.35<sup>a</sup> | 1093±57.98<sup>c</sup> | 1118.5±111.02<sup>b</sup> |
|  | Pt (min) | 9.15±0.35<sup>a</sup> | 8.7±0.35<sup>a</sup> | 8.7±0.42<sup>a</sup> | 9.1±0.42<sup>a</sup> |
|  | PT (°c) | 62.88±0.17<sup>a</sup> | 56.23±1.09<sup>b</sup> | 63.38±3.0<sup>b</sup> | 62.525±2.09<sup>b</sup> |
| Qulle | PV (cp) | 7832±29.70<sup>a</sup> | 7085.5±36.06<sup>b</sup> | 7536.5±17.68<sup>c</sup> | 7509±56.56<sup>d</sup> |
|  | TV (cp) | 2498±33.94<sup>b</sup> | 2265.5±140.71<sup>a</sup> | 2505±26.87<sup>a</sup> | 2495.5±37.48<sup>a</sup> |
|  | FV (cp) | 1307.5±12<sup>c</sup> | 3362±162.6<sup>b</sup> | 3493±60.8a | 3800±74.95<sup>a</sup> |
|  | BV (cp) | 5017.65±13.22<sup>b</sup> | 4542±216.37<sup>a</sup> | 5233.5±129.40<sup>a</sup> | 4984±60.81<sup>b</sup> |
|  | SV (cp) | 2341±2.83<sup>a</sup> | 1061.5±71.42<sup>d</sup> | 920±62.22<sup>c</sup> | 1245.5±28.99<sup>b</sup> |
|  | P t (min) | 6.95±1.35<sup>a</sup> | 8.95±0.07<sup>a</sup> | 8.9±0.14<sup>a</sup> | 8.3±0.56<sup>a</sup> |
|  | PT (°c) | 73.07±2.736<sup>a</sup> | 60.6±1.98<sup>b</sup> | 59.55±0.64<sup>b</sup> | 61.975±1.45<sup>b</sup> |

Values are means duplicates ± SD; Means not sharing a common superscript letter across the rows were significantly different (P<0.05).
Note: FSND: Fermented Sundried, FFBD: Fermented Fluid Bed Dried, FFD: Fermented & freeze dried.

Setback viscosity (SV) can be considered a measure of tendency for flour or starch to retrograde, due to realignment of amylose molecules. The index of retro gradation during cooling is known as setback viscosity. Retro gradation describes the process in which a heated starch paste cools, and the exuded amylose molecules re-associate and unite the swollen starch grains in an ordered structure that results in viscosity increase [44]. Set back viscosity in this study was significantly (P<0.05) increased from 258.5 to 2341cp. The SV of cassava flour in this study was significantly affected by both processing and varieties. The findings of this study show that processed cassava flour has a higher tendency to retrograde than raw cassava flour.

FV is characteristic for the final product quality of starch-based foods [40]. FV of cassava flour in this study was significantly increased from 1307cp in raw qulle varieties to 3800cp in FFD qulle varieties respectively and the results were close to that reported by [43] in which the FV of cassava flour was 2514.67cp. The significant difference (P<0.05) was shown for both varieties and all processing conditions in terms of final viscosity. Increase of FV might be associated with activities of amylases that hydrolyses amylopectin and thus releases polymeric chains of monomer units [45].

Pasting temperature represent the temperature at which the sample will cook. The variation in pasting temperature of the flour could be due to differences in the granules size [44]. Lower pasting temperature and rapid rise in peak viscosity indicates a weak granular structure [42]. The pasting temperature of cassava flour in this study (56-73°C) were close to the values previously reported for cassava flour by [44] in which the pasting temperature of cassava flour was between (60-80°C). According to the finding we suggest that cassava root flour has better Visco-elastic properties and can be best ingredient for bakery recipes

### 2.12. Vitamin C and total carotenoid content of cassava root and leaves

Table 6, shows Vitamin C and Total carotenoid contents for the different cassava root and leave cultivars. The vitamin C content of cassava root was ranged between 5.6 and 19.4%. The smallest value (5.6%) observed for FSND and the highest value (19.4%) was recorded in raw kello variety. The values were in the range between that reported by [46] in which the Vitamin C content of cassava root was between 1 to 40%. The values recorded were significantly (p<0.05) affected by both varieties and processing conditions. This indicates that vitamin C content in cassava roots might be decreased with different processing conditions like fermentation and drying and genetic variations[47]. Whereas, the vitamin C contents of cassava leave were in the range between 1.3 and 29.5%. There was a significant difference (p<0.05) in vitamin C contents for varieties, processing and their interaction. According to [19], vitamin C content of cassava leaves were in the range between 28.3 and 431.5mg/100g. The values in this study were greater than that reported by [48], in which the vitamin C contents of cassava leaves were in the range between (55.07 and 270.23 mg/100g). The difference might be due to the fact that the destruction of Vitamin C during processing such as fermentation.

**Table 6:** Vitamin C and Total carotenoid content of cassava root and leaves as affected by varieties and processing.

| Cassava root |  | Cassava leaves |  |  |  |  |  |  |  |
| --- | --- | --- | --- | --- | --- | --- | --- | --- | --- |
| Varieties | Parameters | Types of processing |  |  |  | Types of processing |  |  |  |
|  |  | Raw | FSND | FFBD | FFD | Raw | BUFBD | BFFBD | BFFD |
| Kello | TCC($\mu\text{g/g}$) | 37.39±0.18<sup>b</sup> | 40.49±0.90<sup>b</sup> | 44.37±1.91<sup>b</sup> | 46.28±1.38<sup>a</sup> | 654.44 ±11.99<sup>c</sup> | 234.10±6.85<sup>d</sup> | 715.99 ±32.59<sup>b</sup> | 827.26±23.60<sup>a</sup> |
| | Vitamin C (mg/100g) | 19.4±1.37<sup>a</sup> | 5.6±0.08<sup>c</sup> | 6.66±0.09<sup>b</sup><sup>c</sup> | 7.53±0.09<sup>c</sup> | 29.144 ±2.69<sup>a</sup> | 2.996 ±0.28<sup>b</sup> | 1.33 ±0.43<sup>b</sup> | 1.545 ±0.07<sup>b</sup> |
| Qulle | TCC($\mu\text{g/g}$) | 31.73±0.52<sup>b</sup> | 37.28±6.35<sup>b</sup> | 33.09±2.7<sup>b</sup> | 49.95±0.08<sup>a</sup> | 325.13±10.33<sup>c</sup> | 239.97±20.62<sup>d</sup> | 819.91±11.88<sup>b</sup> | 987.95±46.29<sup>a</sup> |
| | Vitamin C (mg/100g) | 14.28±0.28<sup>a</sup> | 5.91±0.88<sup>c</sup> | 6.22±0.88<sup>b</sup><sup>c</sup> | 7.26±0.28<sup>c</sup> | 19.578 ±1.53<sup>a</sup> | 2.7435 ±0.07<sup>b</sup> | 1.31 ±0.26<sup>b</sup> | 2.26 ±0.32<sup>b</sup> |

Values are means duplicates ± SD; Means not sharing a common superscript letter across the column were significantly different ( $P \leq 0.05$ )
Note TCC: Total Carotenoid Contents, FSND: Fermented Sun Dried and FFBD: Fermented Fluid Bed Dried and FFD: Fermented Freeze Dried, BUFBD: Boiled Unfermented and fluid bed dried, BFFBD: Boiled Fermented Fluid Bed Dried and BFFD: Boiled Fermented Freeze Dried Freeze Dried

The total carotenoid content of cassava root flour was ranged between 31.73 and 49.95μg/g. The values were significantly (p<0.05) greater in FFD qulle variety as compared to raw cassava root flour. There was significant difference (p<0.05) in all processing techniques and better carotenoid retention was observed in FFD for both varieties. The total carotenoid losses during heat treatment could be due to carotenoid isomerization and oxidation, which is the breakdown of trans-carotenoid to their cis-isomers. The difference in total carotenoid content may be as a result of one or the combinations of factors, such as heat, light, oxygen and enzymes which can lead to major or minor losses of carotenoids in cassava root during processing [49]. Whereas, the total carotenoid content in two local varieties of cassava leaves were in the range between 234.10 and 987.95μg/g. According to the findings in this study the total carotenoids content (234.10 and 987.95μg/g) was slightly greater than the results reported by [26], in which the total carotene content of cassava leaves ranged between 174 and 547μg/g. For both varieties of cassava leaves, the total carotenoid content of cassava leaves were significantly (p<0.05) affected by methods of processing and types of varieties. Although cassava leaves are detoxified with a lot of water, essential nutrients may be leached during processing; however, fermentation enhances the average carotenoid content of the cassava leaves. A possible reason could be that, as major compositions of cassava (carbohydrates, moisture, and fiber) reduced by hydrolysis during fermentation, the proportion of other minor compositions such as carotenoids will apparently increase [40]. According to [19], the β-carotene content of cassava leaves was in the range between 3.0 41.6 43.6 the results were slightly lower than the values recorded in this study. Because β-carotene is included in total carotenoids, this could be the case.

### 2.13. Anti-nutritional factors in cassava root and leaves

The ant nutrient content of cassava leaves were presented in table 7, Tannins are polyphenols with a relatively large molecular weight (up to 20,000 Da), and their presence in cassava leaves is believed to play a significant role in the low net protein utilization.

**Table 7:**
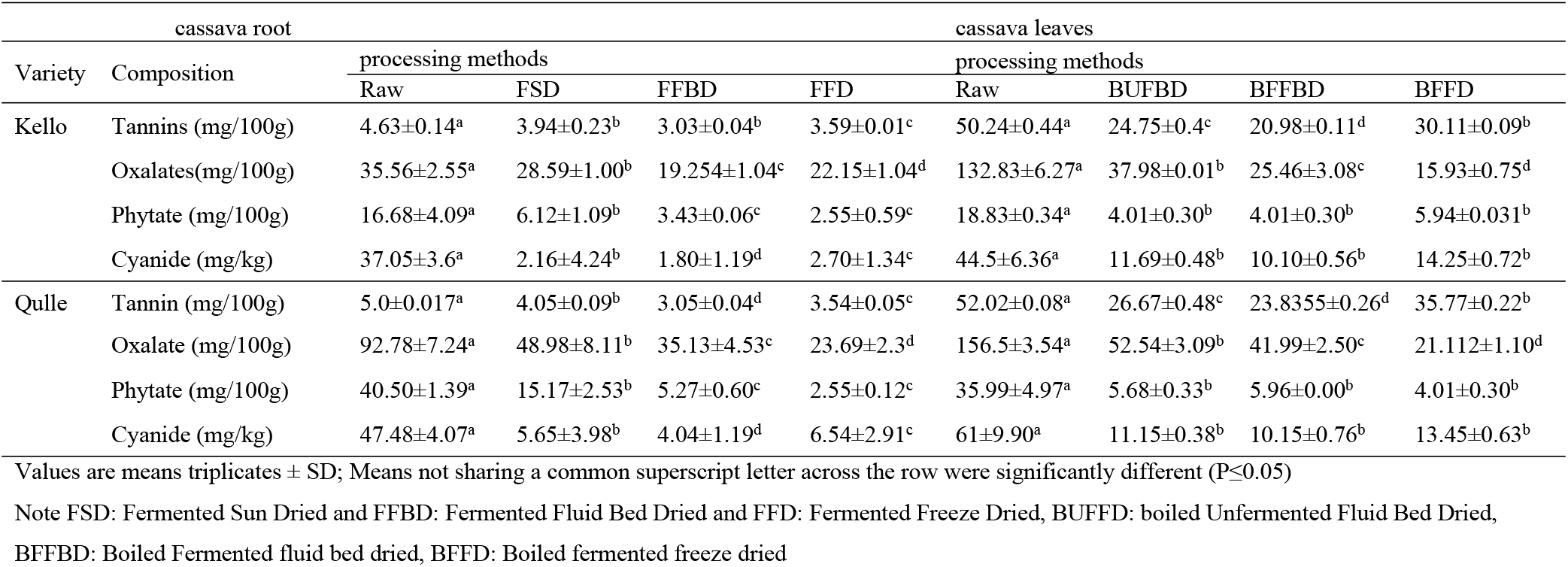
Concentrations of Anti-nutritional factors after processing in two varieties of cassava root and leaves.

| cassava root |  | cassava leaves |  |  |  |  |  |  |  |
| --- | --- | --- | --- | --- | --- | --- | --- | --- | --- |
| Variety | Composition | processing methods |  |  |  | processing methods |  |  |  |
|  |  | Raw | FSD | FFBD | FFD | Raw | BUFBD | BFFBD | BFFD |
| Kello | Tannins (mg/100g) | 4.63±0.14<sup>a</sup> | 3.94±0.23<sup>b</sup> | 3.03±0.04<sup>b</sup> | 3.59±0.01<sup>c</sup> | 50.24±0.44<sup>a</sup> | 24.75±0.4<sup>c</sup> | 20.98±0.11<sup>d</sup> | 30.11±0.09<sup>b</sup> |
|  | Oxalates(mg/100g) | 35.56±2.55<sup>a</sup> | 28.59±1.00<sup>b</sup> | 19.254±1.04<sup>c</sup> | 22.15±1.04<sup>d</sup> | 132.83±6.27<sup>a</sup> | 37.98±0.01<sup>b</sup> | 25.46±3.08<sup>c</sup> | 15.93±0.75<sup>d</sup> |
|  | Phytate (mg/100g) | 16.68±4.09<sup>a</sup> | 6.12±1.09<sup>b</sup> | 3.43±0.06<sup>c</sup> | 2.55±0.59<sup>c</sup> | 18.83±0.34<sup>a</sup> | 4.01±0.30<sup>b</sup> | 4.01±0.30<sup>b</sup> | 5.94±0.031<sup>b</sup> |
|  | Cyanide (mg/kg) | 37.05±3.6<sup>a</sup> | 2.16±4.24<sup>b</sup> | 1.80±1.19<sup>d</sup> | 2.70±1.34<sup>c</sup> | 44.5±6.36<sup>a</sup> | 11.69±0.48<sup>b</sup> | 10.10±0.56<sup>b</sup> | 14.25±0.72<sup>b</sup> |
| Qulle | Tannin (mg/100g) | 5.0±0.017<sup>a</sup> | 4.05±0.09<sup>b</sup> | 3.05±0.04<sup>d</sup> | 3.54±0.05<sup>c</sup> | 52.02±0.08<sup>a</sup> | 26.67±0.48<sup>c</sup> | 23.8355±0.26<sup>d</sup> | 35.77±0.22<sup>b</sup> |
|  | Oxalate (mg/100g) | 92.78±7.24<sup>a</sup> | 48.98±8.11<sup>b</sup> | 35.13±4.53<sup>c</sup> | 23.69±2.3<sup>d</sup> | 156.5±3.54<sup>a</sup> | 52.54±3.09<sup>b</sup> | 41.99±2.50<sup>c</sup> | 21.112±1.10<sup>d</sup> |
|  | Phytate (mg/100g) | 40.50±1.39<sup>a</sup> | 15.17±2.53<sup>b</sup> | 5.27±0.60<sup>c</sup> | 2.55±0.12<sup>c</sup> | 35.99±4.97<sup>a</sup> | 5.68±0.33<sup>b</sup> | 5.96±0.00<sup>b</sup> | 4.01±0.30<sup>b</sup> |
|  | Cyanide (mg/kg) | 47.48±4.07<sup>a</sup> | 5.65±3.98<sup>b</sup> | 4.04±1.19<sup>d</sup> | 6.54±2.91<sup>c</sup> | 61±9.90<sup>a</sup> | 11.15±0.38<sup>b</sup> | 10.15±0.76<sup>b</sup> | 13.45±0.63<sup>b</sup> |

Values are means triplicates ± SD; Means not sharing a common superscript letter across the row were significantly different ( $P \leq 0.05$ )
Note FSD: Fermented Sun Dried and FFBD: Fermented Fluid Bed Dried and FFD: Fermented Freeze Dried, BUFBD: boiled Unfermented Fluid Bed Dried, BFFBD: Boiled Fermented fluid bed dried, BFFD: Boiled fermented freeze dried

They have the potential to create insoluble complexes with proteins, inhibiting digestion by inactivating enzymes. When present in large quantities in food, they can have unfavorable impacts [50]. Thus, the tannin content of cassava root flour in this study was in the range between 3.03 to 5.0mg/100g. A maximum value (5.0mg/100g) was observed in raw kello variety. The contents were significantly (p<0.05) decreased under different processing conditions and also the values were significantly different for different varieties. The level of tannins in this study was close to the literature and the slightly greater than that reported by [48] in which the tannin contents of cassava root were in the range between 2.04 to 4.31mg/100g. The difference may be due to the difference in varietal difference, geographical location, altitude and climatic condition including soil type. Whereas, the tannin contents of cassava leaves as described in table 6, in this study were varied in the range between 20.98 and 52.02mg/100g. The contents observed were significantly different (p<0.05) for both varieties and all processing conditions. [51] reported that the tannin concentration in different varieties of cassava leaves ranged between 6.9 and 15.0mg/100g. According to [21], tannin content of different cassava leaves varieties were in the range between 5.74 and 10.16mg/100g. The values were slightly greater than that reported by literatures. This variation was probably due to the genetic differences among the cultivars, the plant ages, the leaf maturity and soil fertility and processing efficiency.

Oxalates are ant nutrients that negatively affect calcium and magnesium bioavailability. It has the ability to bind calcium and expel it in the urine or form crystals. Because calcium oxalate crystals are a primary component of kidney stones, it is recommended that persons who are prone to them increase their calcium consumption while decrease their oxalate intake [52]. Oxalate content of cassava root flour in the present study was recorded in the range between 22.15 and 92.78mg/100g and the maximum value observed for raw kello and variety (92.78). The oxalate content was significantly differently (p<0.05) affected by processing methods and varieties and also by the combined effect. According to the finding [17], the oxalate contents of the same varieties was reported in the range 24.93mg/100g and 86.18mg/100g for qulle and kello varieties respectively. This indicated that the present result 22.15 to 92.78mg/100g lies within this range of oxalate content of cassava root. But the level of oxalate for raw kello variety was slightly greater than that reported by [17]. The difference in the oxalate level between the experimental value and the reported value maybe as a result of an agro-ecological difference like temperature, climate, soil type, the difference in the performance of the analysts (chemicals) used while performing tests and even the difference in analyst who performed the tests. Whereas, the oxalate content of cassava leaves in this study was ranged between 25.46 and156.5mg/100g. The oxalate contents were significantly (p<0.05) decreased by processing parameters and the values were varied for different varieties. The results obtained were much lower than the results reported by [53] in which the oxalate concentrations of cassava leaves were in range between 1350 to 2880 mg/100g. This indicates the varieties and processing methods used in present study were better in lowering the amount of oxalate.

Phytate is ant-nutritional factor that binds proteins and minerals in the gastrointestinal tract preventing absorption and utilization by the body. Specifically, phytate interferes with the absorption of divalent metals, such as iron and zinc, which are essential nutrients [54]. It has negative impact on the activity of digestive enzymes and act through chelation of mineral cofactors or interaction with protein. For instance, it interferes with zinc homeostasis. Heat processing such as boiling and cooking have no effect in reducing the level of phytic acid as the phytate is relatively heat stable but fermentation does [50]. The phytate content of the two local varieties of cassava root flour in this study was ranged between 2.55 and 40.5mg/100g. The contents were significantly (p<0.05) affected by both processing and varieties. The difference might be due fermentation and intrinsic parameters of cassava leaves varieties. Whereas, the phytate contents of cassava leaves presented in this study were between 4.02 and 36.0mg/100g. The values were significantly (p<0.05) affected by both varieties and all processing techniques. The maximum value (36.0mg/100g) was observed in raw kello variety and the phytate content was varied in different processing methods. This is due to the fact that during cassava leaves fermentation, there might be de-phosphorylation and release of the minerals and makes them available for absorption. Therefore, cassava leaves fermentation minimizes the ant nutritional effect of phytate.

Consumption of 50 to 100mg of cyanide is acute, poisonous and lethal to adults. Lower consumption of cyanide is not lethal but long term intake can cause severe health problem like tropical neuropathy [4]. The total cyanides content of the cassava root flour decreases with processing such as fermentation [42] and thermal treatment [55]. According to results achieved in this study, the cyanide content of the cassava root flour was ranged between 1.8 to 47.48ppm. And the maximum value observed for qulle variety (47.48ppm). The content was significantly (p<0.05) affected by both processing and variety. This indicates that acceptable level of cyanide in cassava root can be achieved by using appropriate processing and by breeding for improved varieties. In addition to the above mentioned processing methods, literature has shown that preprocessing steps such as peeling, grating, and washing significantly reduce the cyanide level of cassava root. According to the reports [36]and [26], cyanide contents of fresh cassava root were ranged between 86.2 to 154.8mg/100g and 346 to 748.4mg/100g respectively. According to the current codex and international standards, the level of ‘total hydrocyanic acid’ in the fresh cassava root flour must not exceed 10 mg kg-1, (FAO/WHO, 2019). The cyanide level of cassava root flour (1.8 to 47.48ppm) in the current study was lower than previously reported levels and was within acceptable international standards. Whereas, the cyanide content of cassava leaves in this study was in the range between 10.10 and 61.0 ppm. And the maximum value (61ppm) was observed in raw qulle variety and the contents were significantly (p<0.05) decreased for various processing techniques and varieties. According to the findings, overall cyanide concentrations decreased to acceptable level on processing with the exception of freeze dried samples. HCN levels were higher in freeze-dried leaves than in other processing methods. This might be due to moisture and nutrient retention capacity of freeze dried food. According to [21] cyanide content of different cassava leaves varieties were in the range between 11.29 and 19.29mg/100g. As per [4], the cyanide level in cassava leaves ranges from 53 to 1300ppm. HCN contents of cassava leaves in this study were close to the results presented by [56] in which the values ranged between (58.5 to 86.7mg/100g). According to [24], the total HCN content found in the cassava leaves varied from 90.6 to 560.9ppm. It’s possible that the little deviation from the standard might be due to inefficient machinery and operations. The current investigation’s cyanide levels (10.10 to 14.25ppm) in different processing methods were found to be below the acute hazardous limit. It is, however, somewhat higher than the WHO’s recommended level of cyanide in final food items, which is 10ppm (WHO/FAO, 2019).

## 3. CONCLUSION

Cassava is a climate-smart, nutrient-dense crop that could be a promising food for the future. In terms of nutritional, physicochemical, and functional qualities, cassava root and leaf flour performed well. Cassava flour’s nutritional and anti-nutritional properties are influenced by the variety and processing method. Despite the fact that cassava is an important food source in developing countries, HCN poisoning is still a problem. Finding appropriate processing that can significantly reduce HCN will be critical in encouraging utilization. The efficiency of these methods is dependent on the method’s nutrition retention and detoxifying capabilities. The concentration of cyanide was found to have decreased in all processing methods, with a minor increase in freeze dried samples, according to the data. This suggests that, despite its superior nutrient retention, freeze drying is ineffective in terms of detoxification. This study confirms that different varieties and a combination of processing methods, such as fermentation, boiling, and drying, modify quality parameters and reduce toxic components significantly (p<0.05).

## ACKNOWLEDGEMENT

The authors are grateful to Mr. Asefa Cholo for his cooperation during sample collection, Mr. Berihun Alebel for his assistance on laboratory analyses, and Mr. Zemanu Kerie for his cooperation on paper work. We also thank Areca Agricultural Research Center in Ethiopia for providing the cassava root and leaf samples used for the research.

